# Dissecting the molecular landscape of Parkinson’s disease and Parkinson’s disease dementia using highly efficient snRNA-seq (HIF-snRNA-seq)

**DOI:** 10.1101/2025.03.01.640894

**Authors:** Sung-Ung Kang, Jinhee Park, Shinwon Ha, Dongsan Kim, Olga Pletnikova, Javier Redding-Ochoa, Juan C. Troncoso, Quan Peng, Beth O. Van Emburgh, Jaldhir Trivedi, Saurav Brahmachari, Bardia Nezami, Valina L. Dawson, Ted M. Dawson

## Abstract

This study presents a transcriptomic analysis of the cingulate cortex (CING) in Parkinson’s disease (PD) and Parkinson’s disease dementia (PDD) using a High-efficiency single-nucleus RNA sequencing (HiF-snRNA-seq) protocol optimized for post-mortem brain samples. RNA quality prediction, poly-A tailing, and dCas9-targeted depletion enabled analysis of 77 high-quality samples from 240 cases, yielding over 2 million nuclei classified into seven major cell types. Disease conditions revealed altered astrocyte and microglia proportions, implicating their roles in neuroinflammation. Differential expression analysis identified unique and shared genes across PD and PDD, linked to synaptic remodeling, stress responses, and inflammation. Stage-specific analysis uncovered tau-dependent early-stage genes and inflammation-associated late-stage genes. This study highlights the CING’s central role in PD and PDD pathophysiology, offering insights into disease mechanisms and identifying candidate genes and pathways for therapeutic and biomarker development.

## Introduction

Parkinson’s disease (PD) and Parkinson’s disease dementia (PDD) are neurodegenerative disorders that impact the central nervous system, characterized by distinct clinical and pathological features.^1^ While PD is predominantly defined by motor symptoms such as tremors, rigidity, and bradykinesia, PDD encompasses cognitive decline and dementia in addition to motor impairments.^2–4^ The transition from PD to PDD is marked by significant neuropathological changes, including the spread of Lewy body pathology and the presence of Alzheimer’s disease (AD)-related features, such as tau tangles and amyloid-β deposits.^5,6^ This convergence of PD and AD pathology highlights the complexity of the mechanisms underlying neurodegeneration and cognitive deficits in PDD.^7–9^

The cingulate cortex (CING), a brain region integral to cognitive and emotional processes such as attention, memory, and decision-making, plays a critical role in the pathophysiology of both PD and PDD.^10,11^ CING dysfunction has been linked to deficits in executive function and attention, while its atrophy is associated with the cognitive impairments observed in PDD.^12,13^ Moreover, functional imaging studies suggest that hyperactivity in the CING may reflect compensatory mechanisms in response to deficits in other brain regions, particularly during the early stages of disease progression.^12^ These findings underscore the importance of the CING as a central hub for neuroinflammatory processes and protein aggregation in PD and PDD.^8,14,15^

Single-nucleus RNA sequencing (snRNA-seq) has emerged as a powerful tool for studying the cellular and molecular landscapes of complex tissues. By enabling the resolution of transcriptomic profiles at a single-cell level, this method provides unprecedented insights into cell-type-specific changes in neurodegenerative diseases.^16^

The combinatorial indexing split-and-pool method commonly used in snRNA-seq offers several advantages, including reduced technical variability with lower doublet ratio and experimental compatibility with isolated nuclei from tissues. However, analyzing postmortem brain samples presents inherent challenges, such as limited RNA quality due to post-mortem interval (PMI), storage conditions, and disease-related tissue damage.^17,18^ These factors often lead to fragmented RNAs, commonly referred to as damaged RNAs, which can compromise sequencing efficiency and downstream analyses.^19–21^ Furthermore, traditional methods for assessing RNA quality, such as RNA Integrity Number (RIN), are widely used but have limitations when dealing with degraded tissues.^22^ While RIN values are effective for assessing overall RNA integrity, they may fail to capture the quality of nuclei-specific RNA, particularly in cases with prolonged storage or challenging handling conditions, such as rare neurodegenerative brain samples.^22^ Additionally, external factors like freeze-thaw cycles during sample handling can exacerbate RNA degradation, further complicating analyses. Addressing these limitations requires innovative approaches that prioritize the inclusion of high-quality nuclei while minimizing the risks related to cost and technical noise.^14,23^

Here, we introduce and utilize an innovative single-nucleus RNA sequencing method, named as High-efficiency single-nucleus RNA sequencing (HiF-snRNA-seq), to address the challenges of processing post-mortem brain samples. This technique focuses on the selection of samples with RIN between 1 and 6, employs poly-A tailing to enhance RNA readouts, and incorporates dCas9-targeted depletion to minimize sequencing noise and bias. This enables the generation of a comprehensive transcriptomic profile of the CING in PD and PDD. By integrating cell-type-specific and stage-specific molecular changes, it highlights critical pathways involved in PD and PDD, paving the way for developing targeted interventions and pathology-specific treatment strategies.

## Results

From the initial 240 cohorts, a total of 135 patient samples were chosen for single-nucleus RNA sequencing (snRNA-seq), including 45 samples each from control, PD, and PDD cases. Then 77 samples were further filtered and processed for downstream analyses of cell-type specific molecular differences across disease states (Figure 1A). The cohort consisted of 55 males (71.4%) and 22 females (28.6%) (Figure 1B), and Braak stage distribution of samples consisted of stages I (14.3%), II (31.2%), III (32.5%), IV (14.3%), and V (1.3%), while the information of 6.5% samples was not available (Figure 1C). In race distribution, most patients were White (93.5%), followed by Black (3.9%) and Asian (2.6%) (Figure 1D), and ages ranged from 41 to 102 years, with a median age of 78 (Figure 1E).

**Figure 1.**
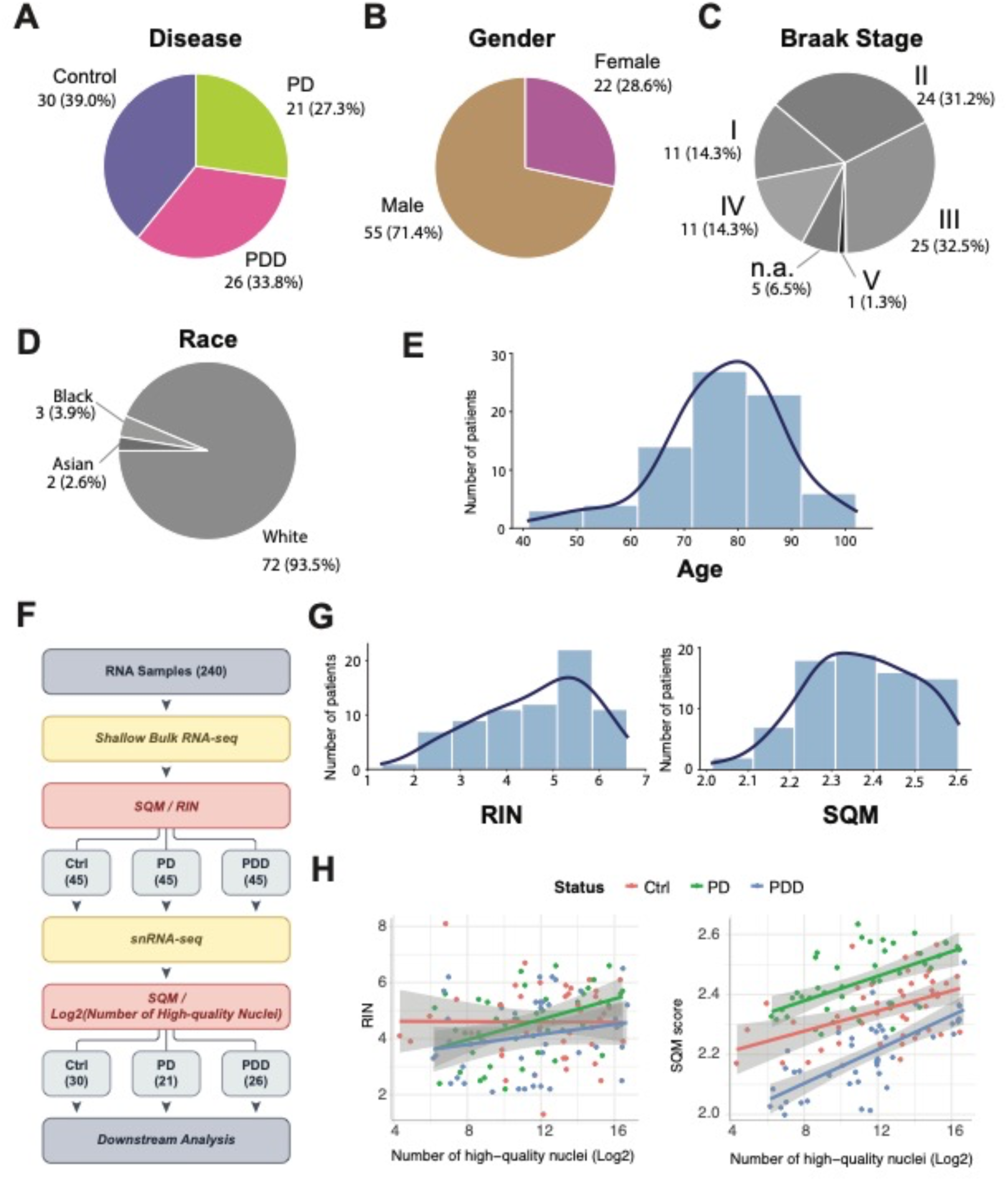
Overview of patient sample selection, demographics, and quality assessment for snRNA-seq analysis. Distribution of the finally selected 77 patient samples by (A) disease, (B) gender, (C) Braak stage, (D) race, and (E) age distribution of the cohort. (F) Illustration depicting the novel SQM metric for sample selection. (G) Comparison of RIN (left) and SQM (right) values across 135 samples. (H) Scatterplots showing correlations between the number of high-quality nuclei and RIN (left) or SQM (right).

### A novel method for estimating sample quality

To select high-quality samples, which have analytical value for snRNA-seq, a quantitative measurement method was developed. This measurement is determined by the genes that are read more or less in high-quality samples, which contain a larger number of high-quality single cells compared to low-quality samples. In this study, a factor, referred to as SQM (sample quality metric), was introduced for the sample selection process (STAR Methods). This approach involved utilizing shallow bulk RNA sequencing across 240 samples to account for molecular diversity, containing reads from high-quality samples (Figure 1F). This approach was specifically designed to address the limitations of traditional methods that rely solely on RNA Integrity Number (RIN) values. This oversimplified RIN-based selection is often inadequate for challenging samples, which are difficult to obtain and often require prolonged storage or unknown post-mortem interval (PMI) such as rare cases, leading to RNA degradation within the tissue. Histograms showed the RIN (Figure 1G, left) and SQM values (Figure 1G, right) for 77 selected samples (30 control, 21 PD, and 26 PDD) out of a total of 240 samples with high measurement values, while ensuring a balanced representation of gender and disease status samples (Figures 1B and 1C). In the comparison of RIN and SQM across groups (control, PD, and PDD), the scatterplots demonstrate a stronger and more consistent positive correlation when SQM values are applied, linking the number of high-quality nuclei with RIN (Figure 1H, left) and SQM (Figure 1H, right) metrics. The results indicate that our measurement using SQM value more accurately reflects sample quality for snRNA-seq compared to the RIN value (Figure 1H). Thus, we employed the SQM values for sample selection and created groups of samples for the multiplexing of the single cell experiment. Each group comprised 15 patient samples, with 5 samples assigned to each specific condition to alleviate batch effects and improve normalization accuracy. Every patient possessed a unique RT-barcode, and we utilized pool and split techniques to combine all the samples (STAR Methods).

### High-efficiency single-nucleus RNA sequencing (HiF-snRNA-seq): Poly-A tailing and dCas9-targeted depletion effectively enhanced snRNA-seq output from post-mortem brain tissue

We employed the widely recognized combinatorial indexing single-cell method to sequence a substantial number of cells extracted from post-mortem brain tissues.^24,25^ However, conventional single-nucleus sequencing methods for these samples are challenging, and numerous researchers have reported difficulties.^19,21^ To address these challenges, we developed an optimized sequencing protocol specifically tailored for post-mortem brain tissue. This protocol incorporates refined methods for poly-A tailing in total RNA-sequencing and dCas9-targeted depletion sequencing, followed by purification, to overcome common issues associated with post-mortem brain samples (Figure 2A).^20,23^ Initially, we applied polyadenylation to capture fragmented RNAs, commonly referred to as damaged RNAs, which may arise due to factors such as postmortem interval (PMI), storage conditions, and disease severity^14^. To account for these factors, RIN values are typically considered. However, a wide range of RIN values introduced biases in selection of samples as well as downstream analyses.

**Figure 2.**
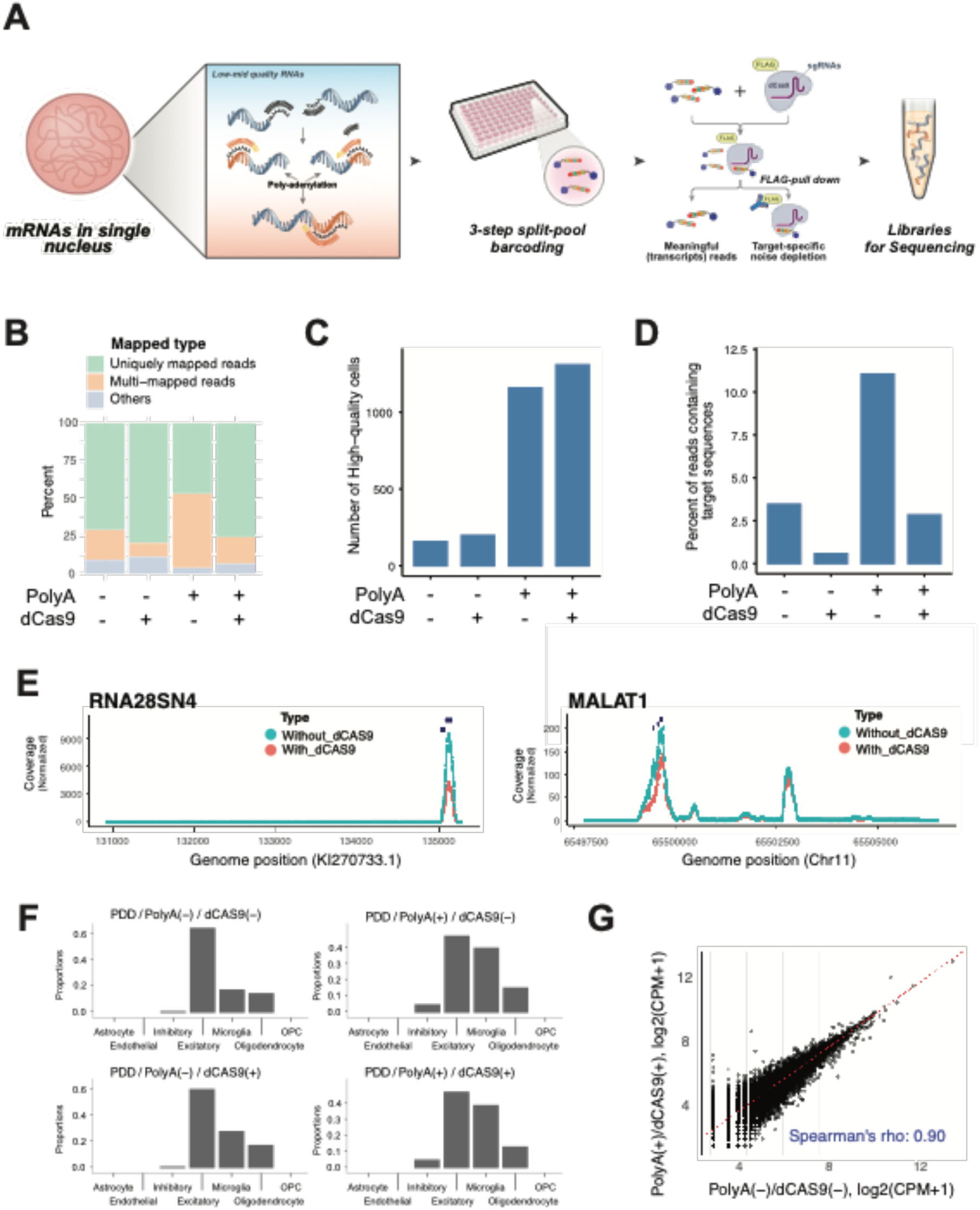
Introduction of HiF-snRNAseq, enhanced sequencing output from post-mortem brain tissues using poly-A tailing and dCas9-targeted depletion. (A) Schematic illustration of the HiF-snRNA-seq procedure. (B) Bar plot for the proportion of uniquely mapped reads, multi-mapped reads, and other non-informative reads across different treatment conditions. (C) Bar plot for the number of high-quality nuclei captured under different treatment conditions. (D) Bar plot illustrating the percentage of reads associated with rRNA content. (E) Coverage plots for RNA28SN4 and MALAT1 regions comparing samples with and without dCas9 treatment. (F) Bar plots showing the proportional representation of major cell types across conditions. (G) Scatterplot showing the correlation of SQM values (log2-transformed) between treated (PolyA+/dCas9+) and untreated samples.

Furthermore, poly-A sequencing results revealed specific artificial poly-A tails in the RNAs within our dataset, prompting the design of dCas9-targeted guide RNAs to enhance mapping of reads (Table S2). These modifications significantly enhance overall sequencing quality (Figures 2B-D). Samples treated with both poly-A and dCas9 showed a higher percentage of uniquely mapped reads than non-treated samples, even after removing a substantial amount of non-informative uniquely mapped reads, such as rRNAs (Figure 2B). This effect subsequently improved the number of high-quality nuclei available for downstream analysis, demonstrating the synergistic effect of the two techniques (Figure 2C). More specifically, the percentage of reads containing target sequences (e.g., rRNA) was significantly reduced when dCas9 was applied (Figure 2D).

As shown in Figure 2E, dCas9 effectively reduced reads associated with specific dCas9-targeted transcriptomic reads (black dots on the coverage plots), including *RNA28SN4* and *MALAT1*, both of which can contribute to sequencing artifacts. To assess the impact of this application on analytical outcomes, snRNA-seq was performed using post-mortem CING tissues from PDD samples. In the aspect on cell type diversity, it showed increased diversity of cell types, particularly the prominence of inhibitory neuron and microglia populations, in conditions such as PDD/PolyA(+)/dCAS9(−) and PDD/PolyA(+)/dCAS9(+) (Figure 2F). Conversely, a scatterplot of log2-transformed CPM (counts per million) values comparing poly- A/dCas9-treated samples to untreated samples demonstrated a strong correlation (Spearman’s rho = 0.90), suggesting no substantial differences in gene expression between the two conditions (Figure 2G). This demonstrates the reliability of the poly- A/dCas9 protocol in preserving biological signal while reducing technical noise.

Therefore, we combined these approaches when using post-mortem brain tissue samples and named it high-efficiency single-nucleus RNA sequencing (HiF-snRNA-seq).

### A 2.0-million-nuclei transcriptional atlas of the human Cingulate Cortex using HiF- snRNA-seq method

To optimize cell type annotation across a total of 2,051,894 nuclei from 77 samples, the average Silhouette coefficient was adopted. The highest Silhouette coefficient was achieved when using SPLIT as the reference source along with an ALRA (Adaptively Learned Rate Adjustment)-adopted principal component dimension, indicating optimal clustering and annotation.^24,26^ The optimal annotation classified the data into seven major cell types including excitatory neurons, inhibitory neurons, oligodendrocytes, oligodendrocyte precursor cells (OPCs), astrocytes, microglia, and endothelial cells, and was visualized by UMAP (Figure 3A, Table S3). The proportions and total numbers of each cell type across the dataset, excitatory neurons were the most abundant, comprising 46.7% of the total cells (957,688 cells), followed by inhibitory neurons (17.1%, 351,676 cells). Oligodendrocytes represented 12.7% (260,776 cells), astrocytes 9.8% (200,790 cells), microglia 8.1% (165,231 cells), endothelial cells 3.4% (70,604 cells), and OPCs accounted for 2.2% (45,130 cells). These data showed a significant representation of neuronal populations while highlighting the presence of astrocyte, glial, and endothelial cells that contribute to the cellular diversity in the CING tissue (Figure 3B). In comparing the cell type proportions across conditions, the overall trends are similar, with excitatory and inhibitory neurons collectively constituting approximately 65% of the total cell population (Figure 3C).

**Figure 3.**
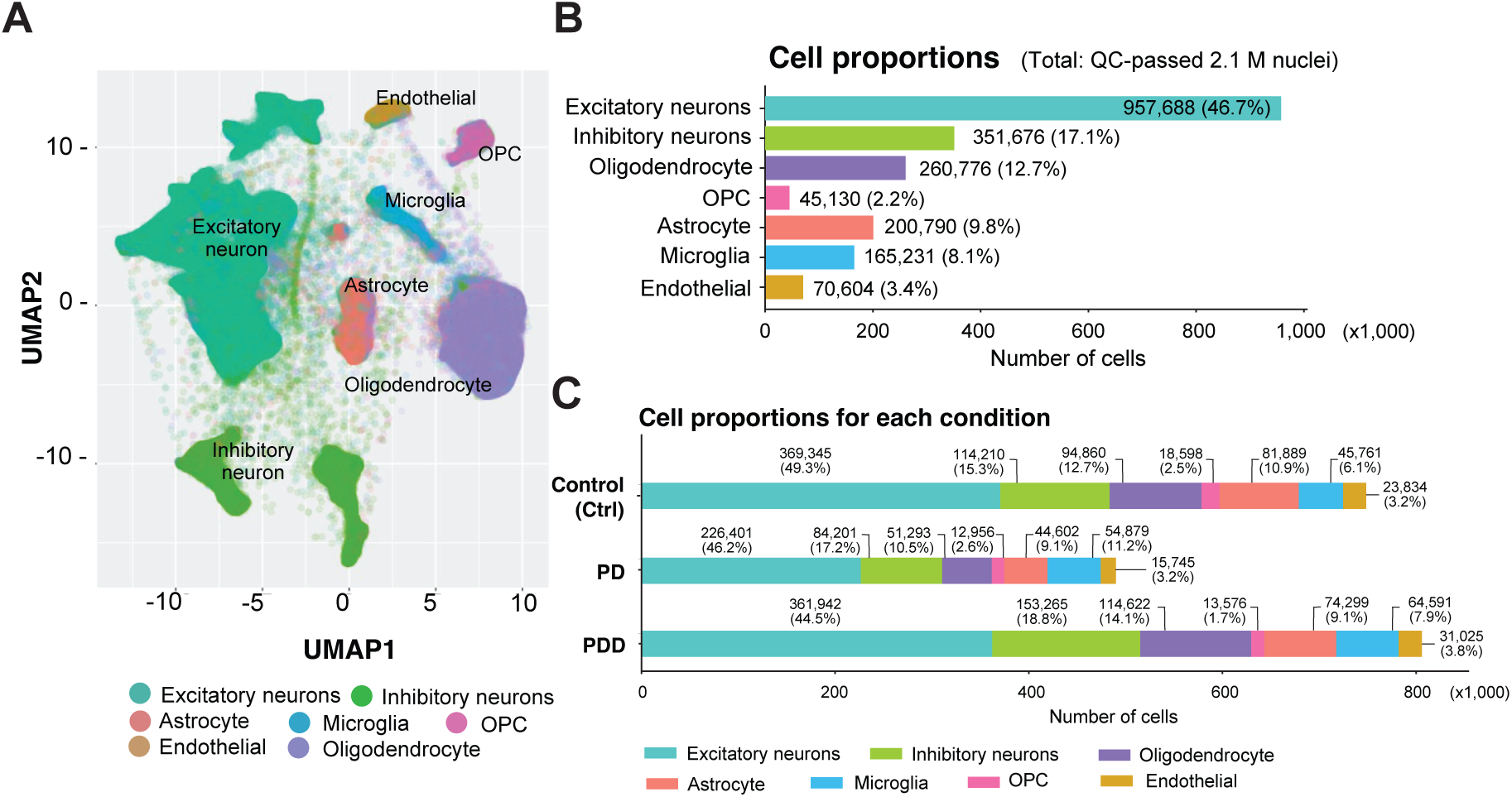
Transcriptional atlas of the human CING in control, PD and PDD. (A) UMAP visualization of 2,051,894 nuclei from 77 samples, classified into seven major cell types: excitatory neurons, inhibitory neurons, oligodendrocytes, OPCs, astrocytes, microglia, and endothelial cells. (B) Bar plot for the total cell proportions of each identified cell type. (C) Bar plot illustrating cell type proportions across three conditions.

Using the chi-square test, we confirmed that there were statistically significant differences in the proportions of all seven cell types. Among all cell types, astrocytes were the only type where the number of cells in both PD and PDD were significantly lower than the expected counts compared to controls. This suggests that transcriptomic changes occur in the astrocyte cell type in both PD and PDD, reflecting both common and disease-specific progression patterns (Figure 3C). These results provide a comprehensive overview of cell type distribution and proportion in both healthy and disease conditions, emphasizing the utility of the optimized annotation pipeline and offering insights into potential cellular mechanisms in PD and PDD.

### Differential gene expression (DGE) analysis in PD, PDD and controls

Differential gene expression (DGE) analysis was performed across datasets: PD versus control, PDD versus control and PDD versus PD, within seven major cell types (Figures 4A-C; Table S4). This analysis aimed to identify both disease-specific and shared differentially expresses genes (DEGs) to elucidate potential functional roles associated with each disease and cell type. In astrocytes, results showed 337 DEGs unique to PD versus control, 495 unique to PDD versus control, and 112 unique to PDD versus PD (Figure 4A). Notably, *CALHM1, CERS1, SLC26A9,* and *PLP1* emerged as unique DEGs in the PD versus controls. These genes are known for involvment in ion homeostasis (*CALHM1*) and sphingolipid metabolism (*CERS1*), processes critical for astrocytic function in neurodegenerative diseases.^27,28^ Additional genes, including *FGD9, GGPS1, CHAC1,* and *HIF1A*, which are linked to stress responses and hypoxia pathways, suggesting astrocyte activation or metabolic stress in PD (Figure 4A; Tables S4 and S5). In the PDD versus controls, representative genes such as *PSEN2, ZIC5, KCNJ8,* and *ALDH1A1* were identified (Figure 4A). *PSEN2* is associated with AD-like pathology^29^, while *ALDH1A1* is known for its role in oxidative stress regulation^30^, highlighting potential astrocytic contributions to PDD pathology. Other genes, including *CD44-AS1, CASP4,* and *CXCL3*, are linked to inflammatory pathways and glial activation, indicating an immune response in PDD.^31–33^ PDD versus PD in CING tissue, revealed distinct DEGs such as *TSSC2, VAMP2, BET1*, and *NPAS4* in astrocytes.

**Figure 4.**
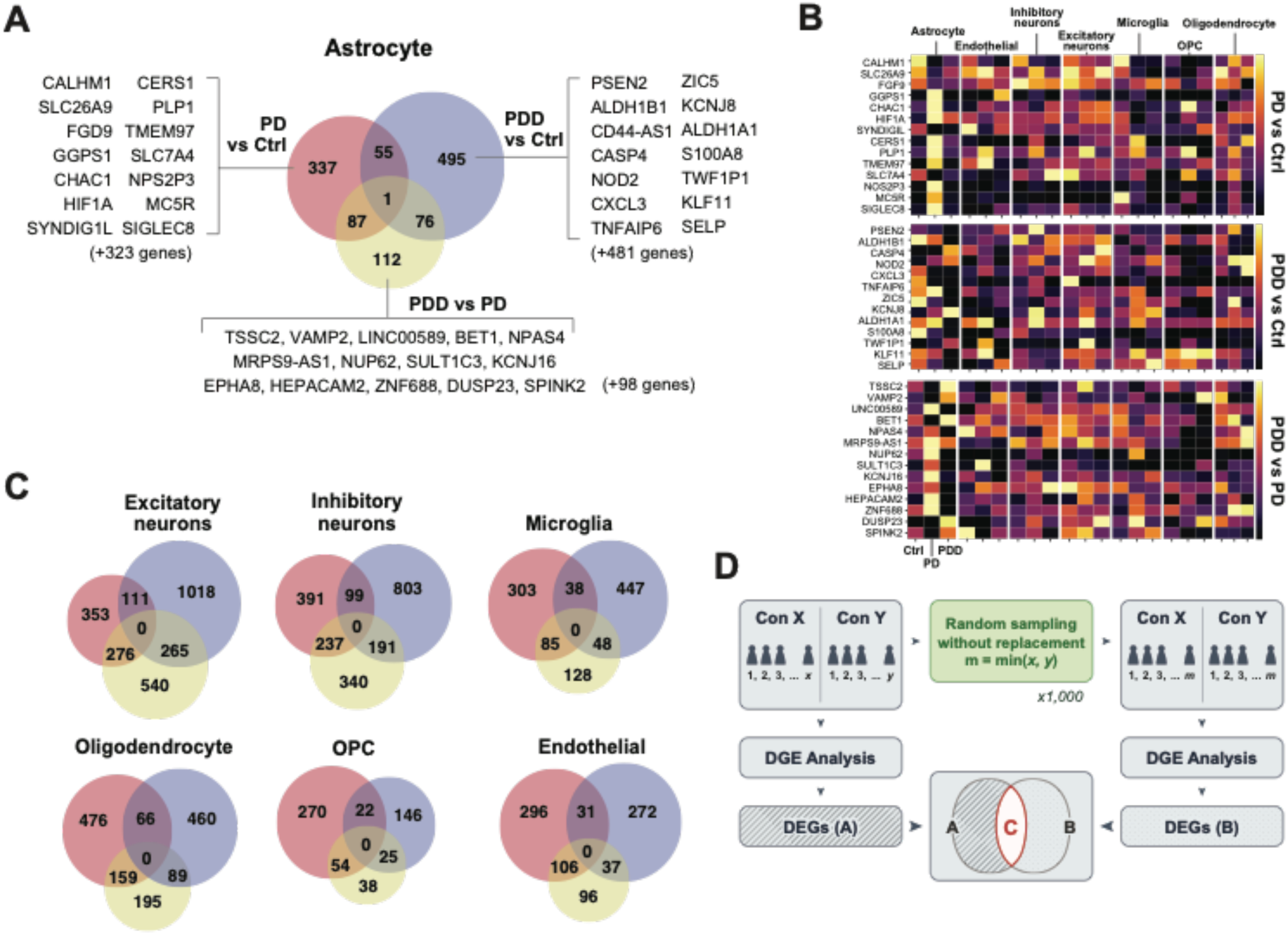
Differential gene expression analysis across PD and PDD conditions in the CING. (A) Venn diagram illustrating differentially expressed genes (DEGs) in astrocytes across three contrasts (PD versus control, PDD versus control, and PDD versus PD) and representative genes. (B) Heatmaps for DEG expression patterns across three contrasts in seven major cell types. (C) Venn diagrams summarizing the number of DEGs in other six cell types across PD and PDD comparisons. (D) Schematic illustration of the permutation test for DEG robustness and stability across cell types.

These genes are associated with synaptic vesicle transport (*VAMP2*) and transcriptional regulation under stress (*NPAS4*), suggesting synaptic remodeling or altered stress responses during the transition to PDD.^34,35^ Additionally, genes like HEPACAM2 and SPINK2 may reflect astrocyte-specific roles in glial signaling and extracellular matrix remodeling during disease progression from PD to PDD (Figure 4A, Table S4).^36,37^ The 55 shared DEGs identified in both the PD versus control and PDD versus control are primarily associated with fundamental stress and inflammatory responses, indicating activation of common pathways in both conditions relative to controls (Figure 4A, Table S4). The shared 87 DEGs identified both in PD versus control and PDD versus PD potentially indicate molecular changes that are initiated in PD and continue to contribute to PDD progression. The shared 76 DEGs identified both in PDD versus control and PDD versus PD represents specific changes to PDD that distinguish it from both PD and control. *TMEM63A*, which is shared across all three comparisons, comprehensively span the spectrum of PD and PDD in astrocyte. A heatmap representation for the representative DEGs across the three comparisons demonstrates clear clustering of disease-specific gene expression patterns (Figure 4B). The DEGs obtained from astrocytes have a specific pattern that are not seen in other cell types. Additionally, we confirmed that the DEGs obtained for all remaining cell types also had cell type specificity (Table S4). This visualization further confirms the distinction between PD and PDD at the molecular level for all cell types. Venn diagrams for the other six cell types shows the number of DEGs specific to PD, PDD and controls and overlapping DEGs among the groups (Figure 4C). These findings support cell-type-specific responses to neurodegenerative processes and further emphasize the unique molecular changes associated with PD and PDD. Using the identified DEGs, KEGG pathway enrichment analysis was conducted and the top 20 enriched pathways for each cell type are provided (Table S5). Notably, pathways such as neuroactive ligand-receptor interaction and complement cascades were enriched in astrocyte DEGs, highlighting their involvement in neuroinflammatory responses (Table S5). In PD, oligodendrocytes showed enrichment in pathways related to dopaminergic synapses and PD, emphasizing their role in PD-specific neurodegenerative mechanisms. In contrast, in PDD, inhibitory neurons and microglia were enriched for AD-related pathways.

Interestingly, microglia exhibited DEGs associated with both AD and PD, indicating a potential dual role in the pathology of PDD (Table S5). These analyses highlight the distinct molecular signatures and pathway enrichments in major cell types across PD and PDD conditions. Permutation tests to provide a measure of the robustness and stability of these analyses were performed using random subsets of samples for each comparison and cell type (1,000 iterations) (Figure 4D, Table 1). This analysis validated the impact of sample size on the DGE analysis and variance between individuals within the same conditions. For each cell type, the number of DEGs identified in the entire dataset (A), the average number of DEGs in random subsets (B), and the number of overlapping DEGs between the two (C) are presented. The average mean and median of Ratio1 (C/A) and Ratio2 (C/B) across cell types are reported for each experimental comparison (Table 1). Ratio 1 represents how the number of samples affects the DGE analysis results, and Ratio 2 indicates the variances between individuals within the same condition. The average median Ratio 1 across cell types in PD versus control, PDD versus control, and PDD versus PD were 0.319, 0.314, and 0.498 respectively, with an overall high robustness index (the average median Ratio 2 median = 0.786 in PD versus control, 0.787 in PDD versus control, and 0.845 in PDD versus PD) signifying substantial overlap across iterations (Table 1). These results showed that comparisons of two disease groups have more similarity than those from the control group. Overall for different cell types and comparisons, approximately 40% of genes are consistently obtained when the sample size was reduced by 20%, and 80% genes are consistently obtained regardless of sample distribution.

**Table 1.**
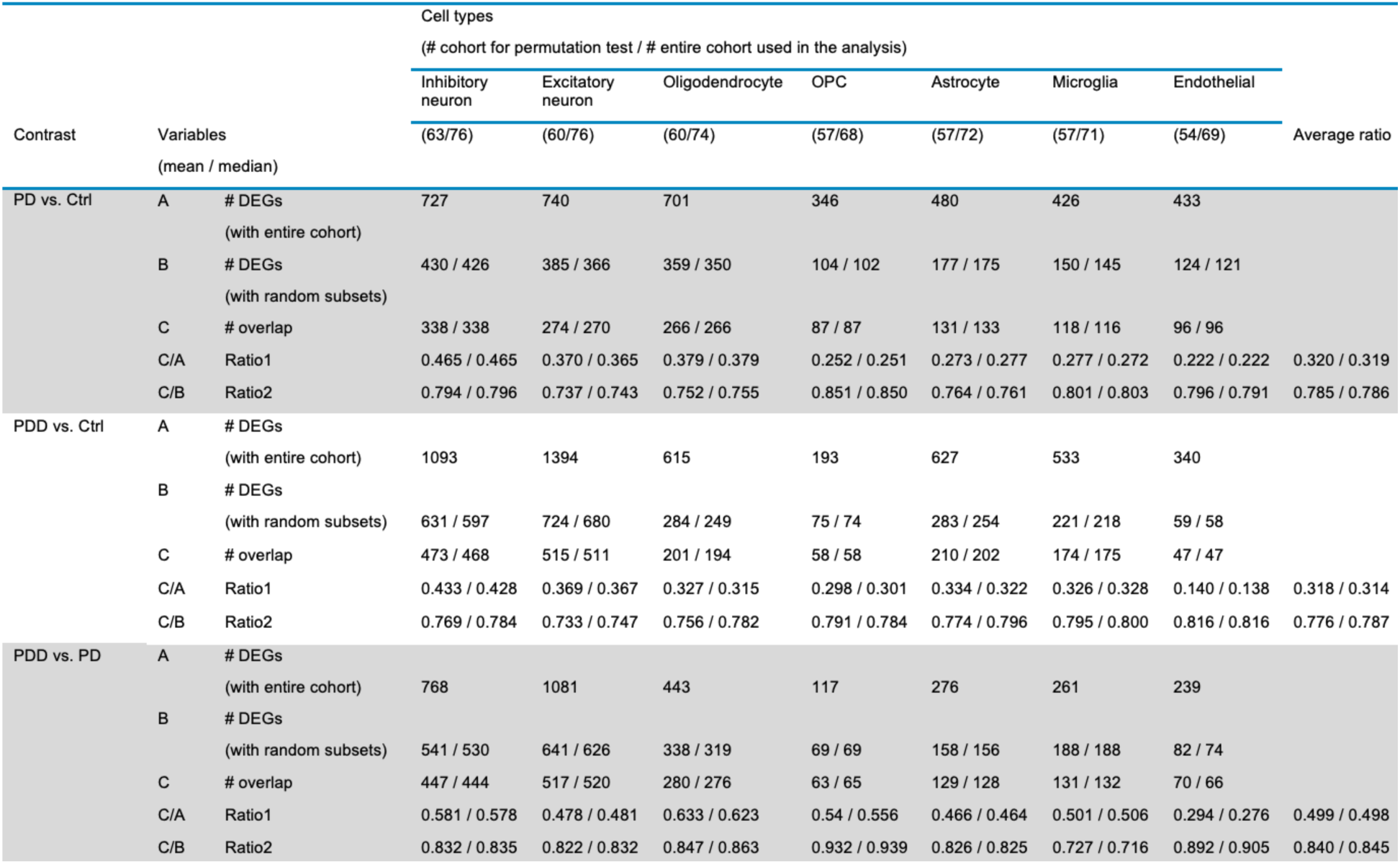
Permutation test results with 1000 iterations using random subsets across experimental contrasts.

| Contrast | Variables<br>(mean / median) | Cell types<br>(# cohort for permutation test / # entire cohort used in the analysis) |  |  |  |  |  |  | Average ratio |
| --- | --- | --- | --- | --- | --- | --- | --- | --- | --- |
|  |  | Inhibitory neuron | Excitatory neuron | Oligodendrocyte | OPC | Astrocyte | Microglia | Endothelial |  |
|  |  | (63/76) | (60/76) | (60/74) | (57/68) | (57/72) | (57/71) | (54/69) |  |
| PD vs. Ctrl | A # DEGs<br>(with entire cohort) | 727 | 740 | 701 | 346 | 480 | 426 | 433 |  |
|  | B # DEGs<br>(with random subsets) | 430 / 426 | 385 / 366 | 359 / 350 | 104 / 102 | 177 / 175 | 150 / 145 | 124 / 121 |  |
|  | C # overlap | 338 / 338 | 274 / 270 | 266 / 266 | 87 / 87 | 131 / 133 | 118 / 116 | 96 / 96 |  |
|  | C/A Ratio1 | 0.465 / 0.465 | 0.370 / 0.365 | 0.379 / 0.379 | 0.252 / 0.251 | 0.273 / 0.277 | 0.277 / 0.272 | 0.222 / 0.222 | 0.320 / 0.319 |
|  | C/B Ratio2 | 0.794 / 0.796 | 0.737 / 0.743 | 0.752 / 0.755 | 0.851 / 0.850 | 0.764 / 0.761 | 0.801 / 0.803 | 0.796 / 0.791 | 0.785 / 0.786 |
| PDD vs. Ctrl | A # DEGs<br>(with entire cohort) | 1093 | 1394 | 615 | 193 | 627 | 533 | 340 |  |
|  | B # DEGs<br>(with random subsets) | 631 / 597 | 724 / 680 | 284 / 249 | 75 / 74 | 283 / 254 | 221 / 218 | 59 / 58 |  |
|  | C # overlap | 473 / 468 | 515 / 511 | 201 / 194 | 58 / 58 | 210 / 202 | 174 / 175 | 47 / 47 |  |
|  | C/A Ratio1 | 0.433 / 0.428 | 0.369 / 0.367 | 0.327 / 0.315 | 0.298 / 0.301 | 0.334 / 0.322 | 0.326 / 0.328 | 0.140 / 0.138 | 0.318 / 0.314 |
|  | C/B Ratio2 | 0.769 / 0.784 | 0.733 / 0.747 | 0.756 / 0.782 | 0.791 / 0.784 | 0.774 / 0.796 | 0.795 / 0.800 | 0.816 / 0.816 | 0.776 / 0.787 |
| PDD vs. PD | A # DEGs<br>(with entire cohort) | 768 | 1081 | 443 | 117 | 276 | 261 | 239 |  |
|  | B # DEGs<br>(with random subsets) | 541 / 530 | 641 / 626 | 338 / 319 | 69 / 69 | 158 / 156 | 188 / 188 | 82 / 74 |  |
|  | C # overlap | 447 / 444 | 517 / 520 | 280 / 276 | 63 / 65 | 129 / 128 | 131 / 132 | 70 / 66 |  |
|  | C/A Ratio1 | 0.581 / 0.578 | 0.478 / 0.481 | 0.633 / 0.623 | 0.54 / 0.556 | 0.466 / 0.464 | 0.501 / 0.506 | 0.294 / 0.276 | 0.499 / 0.498 |
|  | C/B Ratio2 | 0.832 / 0.835 | 0.822 / 0.832 | 0.847 / 0.863 | 0.932 / 0.939 | 0.826 / 0.825 | 0.727 / 0.716 | 0.892 / 0.905 | 0.840 / 0.845 |

### Impact of clinical variables on gene expression: examination of disease progression using Braak stages

Differential expression analysis was performed across seven major brain cell types considering the clinical variables of age, Braak stage, and gender, as well as combinations of these variables (Figure 5A). Among these, Braak stage had the most significant impact on gene expression across all cell types, even when paired with other variables. The number of significant DEGs (FDR < 0.05) was highest in comparisons involving Braak stage, highlighting its central role in driving disease-associated transcriptomic changes (Figure 5A). Additionally, absolute abundances of the seven major cell types were assessed across early (stages I and II) and late (stages III-V) stages. Boxplots summarizing cell abundances revealed no significant changes in the overall composition of these cell types during disease progression, suggesting that transcriptomic rather than compositional changes underlie the observed alterations in disease states (Figure 5B). Stage-specific DEGs were identified for each disease condition across all cell types, and the results highlighted stage-specific transcriptomic signatures (Figures 5C-H, Table S6). Additionally, we confirmed that the DEGs we obtained showed stage- and cell type-specificity in other cell types (Table S6). The overlap of DEGs between early and late Braak stages was minimal across all comparisons, indicating distinct molecular profiles for early and late disease stages (Figures 5C, 5E, and 5G). Early-stage DEGs in the comparisons of PD versus control and PDD versus control were predominantly upregulated, while late-stage DEGs were largely downregulated (Figures 5D and 5F). When comparing PDD versus PD, early-stage DEGs showed higher expression levels in PD than in PDD and late-stage DEGs showed higher expression levels in PDD than in PD (Figure 5H). Because this stage-specific analysis was performed using stratified control samples, an additional experiment using unstratified control samples was conducted (Figure S1, Table S7). In addition, the stage-specific DEGs using stratified control samples were compared with the DEGs obtained by using the entire sample, especially the DEGs unique to each contrast. This analysis revealed a subset of robust candidate genes consistently associated with both disease progression and disease condition-specific contrasts (Figure S2, Table S8). DEGs that are unique to each contrast and overlap with early- or late-stage DEGs are more likely to contain potential biomarker candidates and are suitable for further validation (Table S8). Furthermore, using shallow bulk sequencing data from the entire sample set, which had been previously analyzed as part of quality control, a similar analysis was performed by dividing the data into early and late stages (Figure S3A). This analysis highlighted representative genes, such as *CA2* in microglia, which showed higher expression in PD compared to the stratified control group.

**Figure 5.**
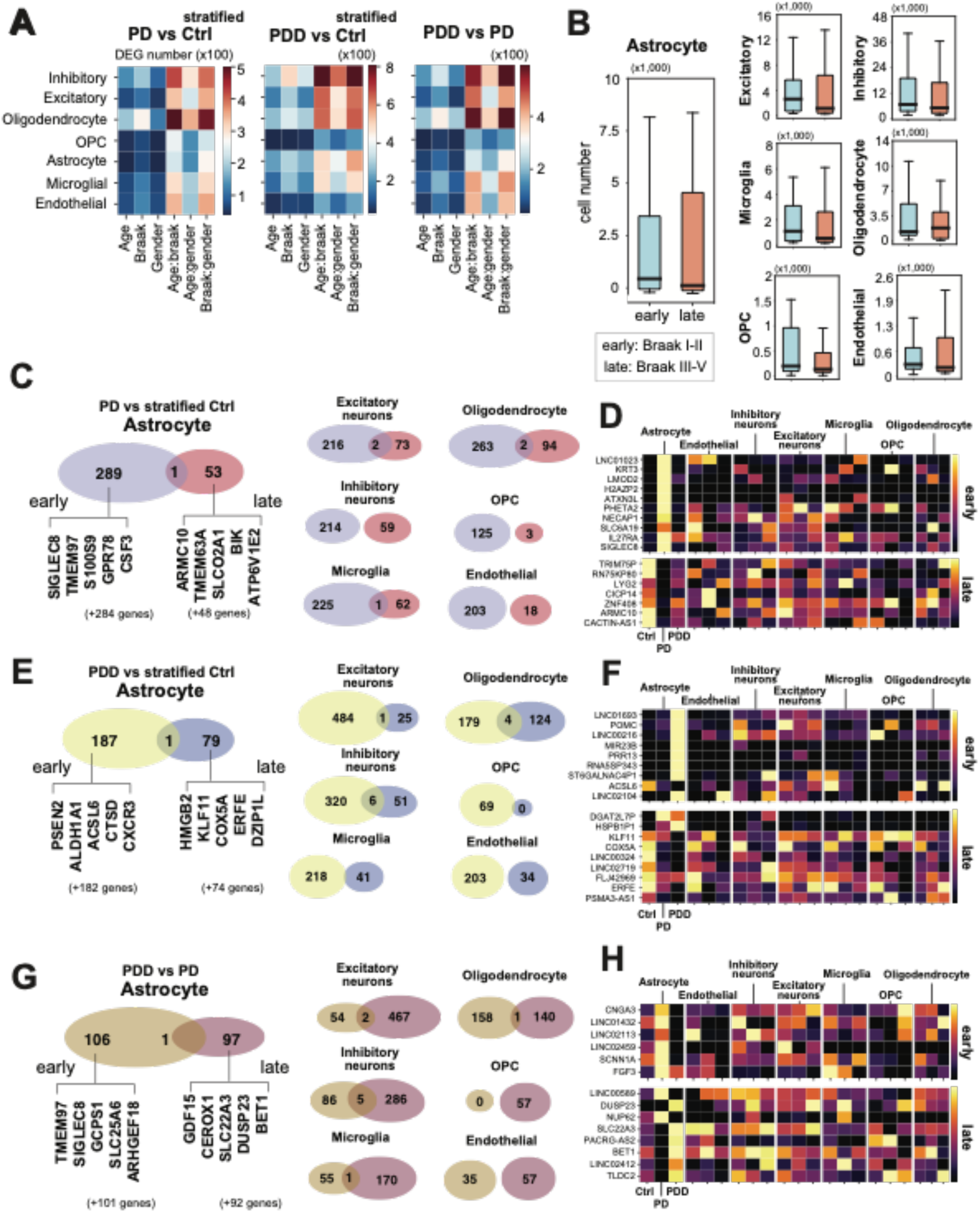
Impact of clinical variables and Braak stage on transcriptomic profiles in PD and PDD. (A) Heatmap presenting the number of significant DEGs (FDR<0.05) across seven cell types (row) and six variables (columns) within three contrasts, PD versus control, PDD versus control, and PDD versus PD. (B) Boxplots representing the absolute abundances of seven cell types across early (Braak stages I-II) and late (Braak stages III-V) stages of disease progression. (C) Venn diagrams displaying the overlap of early- and late-stage DEGs across cell types for the PD versus control comparison. Representative unique genes listed in Astrocyte. (D) Heatmap of representative DEGs for early and late Braak stages in the PD versus control comparison, categorized by sever cell types. (E) Venn diagrams for the PDD versus control comparison, (F) with heatmap of representative DEGs for PDD versus control, and (G) Venn diagrams for the PDD versus PD comparison, (H) with Heatmap representing DEGs in the PDD versus PD comparison.

Furthermore, genes such as *SERPINH1* in astrocytes, *NEAT1* and *DNAJB1* in microglia, *HSPB1* in excitatory neurons, and *NR2F6* in endothelial cells exhibited elevated expression in PDD compared to controls (Figure S3B, Table S9).

### Gene expression insights into Tau-dependent and independent mechanisms in PD and PDD

Post-mortem brain tissues were provided by two brain banks, the Johns Hopkins University School of Medicine and the Banner Health Institution. Notably, these two institutions differ in their methods for assessing Braak staging. At Johns Hopkins Medicine, Braak staging focuses specifically on tau lesions, using neurofibrillary tangles as the primary marker. The progression is strictly estimated across stages I–V, starting in the entorhinal cortex and advancing to the hippocampus, neocortex, and visual cortex. In contrast, the Banner Health Institution employs a more integrated evaluation system that includes tau-based neurofibrillary tangles, CERAD (Consortium to Establish a Registry for Alzheimer’s Disease) scoring for amyloid plaques, and the USSLDB (Unified Staging System for Lewy Body Disorders) to assess α-synuclein pathology.

The core methodology shared by both institutions lies in their evaluation of neurofibrillary tangles and the progression of tau pathology in specific brain regions.^38–40^ Therefore, we thought that analyzing DEGs based on changes in Braak stages, irrespective of the disease, alongside DEGs identified from each specific disease, could provide insights to distinguish between Tau-dependent and disease-related DEGs as well as Tau-independent genes associated with disease progression. Therefore, to further refine the understanding of the molecular mechanisms underlying PD and PDD, we divided all 77 samples solely by Braak stage (early and late) and conducted DGE analysis. These results were compared with previously identified disease stage-specific disease-associated gene sets to discern genes related specifically to Braak stage-associated tau pathology from those linked to disease progression independent of tau pathology (Figure 6A). Early-stage DEGs in astrocytes include 57 overlapping genes, such as *CDC42EP3P1, SH3BP1, GPHB5, FOLR1, BMP10*, and *ALDH3A1*, which are strongly associated with both tau-related pathology and early-stage disease progression (Figure 6B, Table S10). SH3BP1 is known for its role in cytoskeletal organization and neuronal morphology, while ALDH3A1 plays a role in oxidative stress defense and neuroprotection, and FOLR1 is essential for folate transport, critical for neurodevelopment and neuronal maintenance.^41–44^ In contrast, 232 non-overlapping genes, such as *TMEM97, S100A9*, and *SIGLEC8,* were identified as PD-specific genes independent of tau-related changes (Figure 6B, Table S10). *S100A9*, a pro-inflammatory marker, is associated with glial activation and neuroinflammation in PD, while *SIGLEC8*, involved in modulating immune responses, has been implicated in reducing inflammation through glycan recognition.^43,45^ At the late Braak stage, astrocytes exhibited fewer DEGs compared to the early stage, with genes such as *MEIS3P2, CACTIN-AS1,* and *C1QTNF9B* being significantly changed and potentially linked to late-stage glial responses, complement cascade activation, and chronic inflammation.^46,47^ Additionally, 25 non-overlapping genes, such as *ATP2A1* and *miR3609*, were identified as candidates for late-stage PD-specific changes independent of tau pathology. ATP2A1 is known to modulate calcium homeostasis, critical for neuronal signaling.^48^ In PDD, a similar trend with distinct gene sets in early and late Braak stages were observed. Ninety-five overlapping genes in astrocytes during the early stage were identified, including *RHOA-IT1, SLC6A2,* and *CXCR3* (Figure 6C, Table S10). These are associated with tau pathology and early-stage PDD progression, with CXCR3 known to regulate immune cell trafficking and neuroinflammation.^49^ Ninety-two non-overlapping genes, such as *PSEN2, CTSD, GPRIN1*, and *ALDH1A1*, were highlighted as specific to early PDD progression independent of tau pathology. PSEN2 mutations are strongly linked to neurodegenerative processes, particularly in Alzheimer’s and dementia, and ALDH1A1 plays a role in detoxification and has been identified as a marker of astrocytic activity in neurodegeneration.^29,30^ Late-stage PDD was characterized by fewer overlapping genes, such as *MEIS3P2, DGAT2L7P, HSPB1P1,* and *PSMA3-AS1*, and 49 non-overlapping genes (Figure 6C, Table S10).

**Figure 6.**
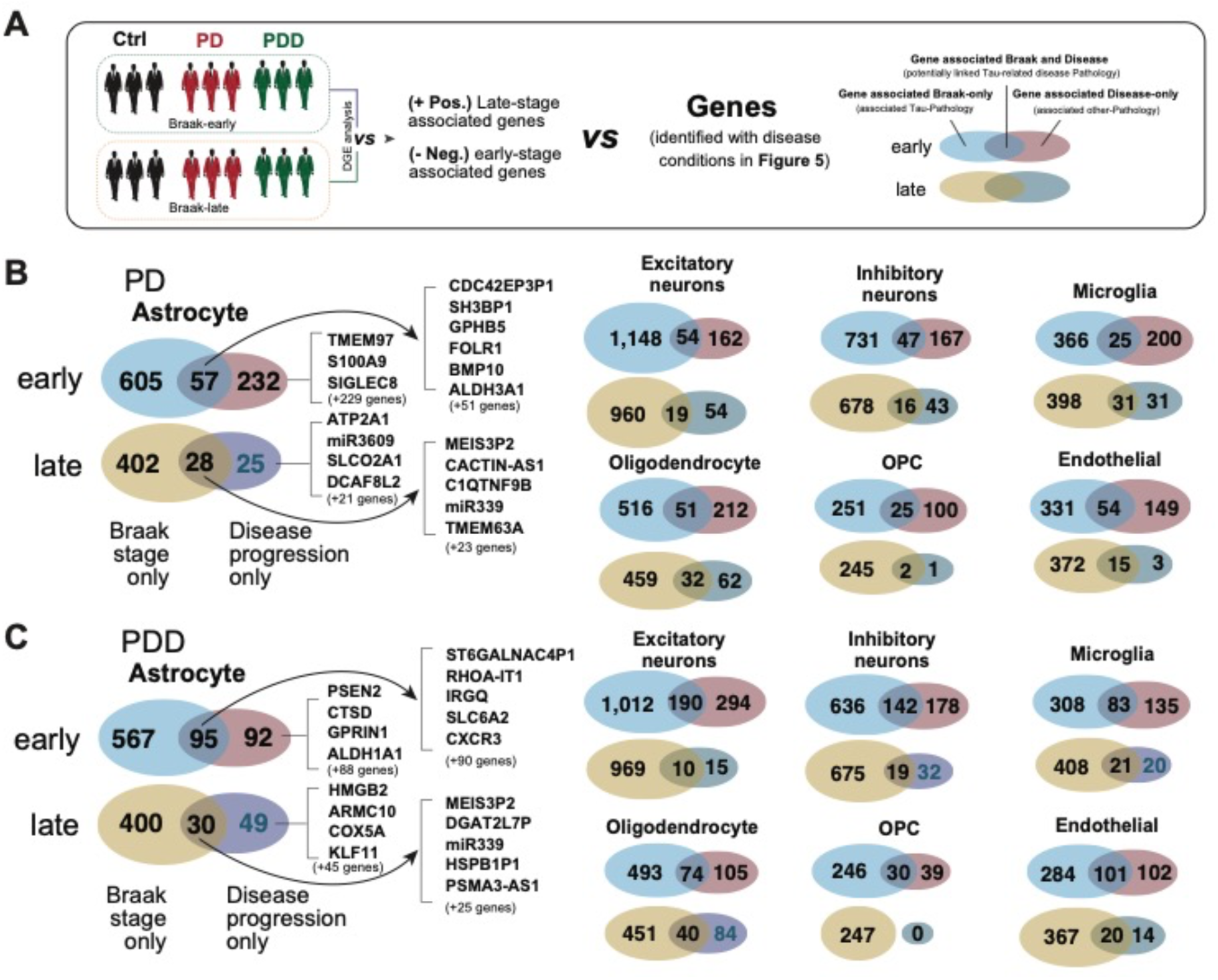
In-depth analysis of disease progression and Braak stages. (A) Workflow illustrating the comparison of genes associated with Braak stage (early and late) and their relationship to tau-related disease progression. The representative genes in astrocyte and the number of DEGs (B) in PD and (C) in PDD across seven cell types are shown.

Overlapping genes are associated with both tau pathology and disease progression, where HSPB1P1 functions as a molecular chaperone involved in protein folding and protection against oxidative stress, which is particularly critical in late-stage neurodegenerative diseases.^50^ In addition to astrocytes, all major brain cell types showed unique DEGs according to cell type and Braak stages (Figure 6). The division of samples based on Braak stage provided a unique perspective to disentangle tau-related pathology from disease-specific mechanisms. By focusing on both Braak-dependent and independent DEGs, this analysis highlights candidate genes and pathways that could serve as targets for further functional validation or therapeutic intervention.

## Discussion

This study provides a comprehensive transcriptomic landscape of the CING in PD and PDD, revealing cell-type-specific molecular changes and stage-specific pathways. The CING is a critical region implicated in the pathophysiology of PD and PDD, serving as a central hub for neuroinflammatory processes and protein aggregation.^7–9,15^ The detection of Lewy bodies and AD-related pathology, such as tau and amyloid-β deposits, within the CING highlights its significant role in the cognitive deficits characteristic of these disorders.^8,13,51^ This convergence of PD and AD-associated neuropathological markers underscores the complex interplay of neurodegenerative mechanisms in the CING, presenting a valuable avenue for exploring both shared and disease-specific pathways underlying motor and cognitive impairments.^13,52^ In the sample selection, the study included a total of 77 patient samples for snRNA-seq, comprising 30 controls, 21 PD cases, and 26 PDD cases, which were carefully selected from an initial cohort of 240 samples (Figures 1A and 1F). The demographic characteristics of the cohort provide critical context for interpreting the molecular findings. The inclusion of samples across a wide range of Braak stages (I-V) provides insights into the molecular transitions from early to advanced disease, highlighting the role of the CING in different phases of neurodegeneration (Figure 1C).

The primary challenge in processing post-mortem brain tissues for snRNA-seq lies in the variability of RNA quality (Figure 1G). While known factors such as the post-mortem interval (PMI) and storage duration significantly influence RNA integrity^14,53^, less well-characterized factors, such as freeze-thaw cycles (transitions between -80°C and - 20°C) during sample handling and distribution, as well as potential contamination, can further exacerbate RNA degradation.^54,55^ These additional factors often lack detailed documentation, making it difficult to match them with initial RNA quality information. This issue is particularly pronounced in rare and high-demand cases such as PD and PDD. Traditional measures of RNA quality, such as RNA Integrity Number (RIN), are widely used but often fail to adequately capture sample quality in degraded tissues. In particular, selecting samples with a RIN value between 2 and 6 for costly snRNA-seq presents a significant challenge for researchers. To address these challenges, this study introduced a quantitative metric, SQM (sample quality metric), to assess sample quality and ensure the inclusion of relatively high-quality samples for snRNA-seq analysis (Figures 1F-H). Scatterplots comparing SQM and RIN values demonstrated a stronger positive correlation for SQM, reflecting its superior ability to accurately predict the suitability of samples for downstream analyses (Figure 1H). This innovative approach is especially useful for analyzing tissue samples with low to mid-range RIN values, as well as long-preserved tissue samples in the rare cases.

The integration of poly-A tailing and dCas9-targeted depletion into the sequencing protocol, named HiF-snRNA-seq, showed a significant advancement in handling post-mortem brain samples (Figure 2). By targeting and removing common artifacts, such as ribosomal RNA (rRNA) and mitochondrial RNA (mtRNA), these steps significantly improved the proportion of uniquely mapped reads and minimized biases introduced by variability in RNA integrity, which is particularly problematic in post-mortem tissues (Figures 2B-E).^20,23,56,57^ Therefore, these advancements are helpful for investigating neurodegenerative diseases such as PD and PDD, where RNA degradation and variability frequently pose significant challenges.

Using HiF-snRNA-seq, we successfully classified over 2 million nuclei from 77 patient samples into seven major cell types (Figure 3A). This cellular atlas of the human CING tissue revealed a significant representation of excitatory neurons (46.7%) and inhibitory neurons (17.1%), underscoring the predominance of neuronal populations in this brain region (Figure 3B). The atlas highlighted that the decreased proportion of astrocytes and the corresponding increased proportion of microglia observed in PD and PDD may represent disease-associated glial responses within the CING region, a hallmark of neuroinflammatory processes (Figure 3C).^8,58^ Conversely, the relatively consistent proportions of OPCs and endothelial cells across conditions suggest their stability under pathological states (Figure 3C).^59,60^ This comprehensive cell-type distribution highlights the dynamic interplay between neurons and glial cells in the progression of PD and PDD, providing insights into potential cellular mechanisms underlying these conditions.

DGE analysis identified unique and shared DEGs across PD and PDD, revealing important molecular insights. Particularly, distinct DEGs, such as *TSSC2* and *NPAS4* in astrocytes in the PDD versus PD comparison, highlight synaptic remodeling and stress response pathways specific to the transition from PD to PDD (Figure 4A).^46,61,62^ These findings emphasize the role of astrocytes in neuroinflammatory responses, synaptic changes, and metabolic stress during disease progression, offering potential therapeutic targets for modulating astrocytic function to mitigate neurodegeneration and cognitive decline.^62,63^ Moreover, shared DEGs between PDD versus control and PDD versus PD (76 genes) in astrocytes represent progressive changes as PDD develops, potentially marking the neurodegenerative trajectory (Figure 4A). Pathway enrichment analysis further highlighted the distinct molecular signatures in major cell types across PD, PDD versus controls (Table S5). In PD, astrocyte DEGs were enriched for pathways related to neuroactive ligand-receptor interactions and ion homeostasis, reflecting their involvement in neuroinflammatory responses. DEGs obtained from oligodendrocyte showed enrichment in dopaminergic synapse and PD-specific pathways, emphasizing their role in neurodegeneration (Table S5). In contrast, inhibitory neurons and microglia in PDD were enriched for AD-related pathways, indicating overlapping mechanisms between PD, PDD, and AD. Permutation testing demonstrated the robustness and stability of DEG identification across experimental comparison. Through permutation testing, it was confirmed that the control group had a larger variance than the disease groups, and that the similarity between the two disease groups was higher than between the control group. This robust methodology underscores the reliability of our DEG datasets and enhances confidence in the identified molecular targets for further validation.

Transcriptomic changes in PD and PDD were analyzed using two distinct control groups, a stratified control group and an unstratified control group (Figures 5 and S1). The unstratified control group increased statistical power and emphasized disease-specific changes by utilizing all control samples, while the stratified approach provided higher biological resolution, captured early pathological changes, and ensured Braak stage-matched comparisons. Overlapping genes identified between condition-specific and stage-specific represent markers of disease progression, reflecting core molecular changes, enhancing biomarker reliability and guiding therapeutic development. Cross-referencing these results, therefore, offers a comprehensive understanding of PD- and PDD-related transcriptomic changes and identifies reliable markers for progression and intervention.

Braak stage-specific DGE analysis provided critical insights into tau-dependent and independent molecular changes. Early-stage DEGs in astrocytes, such as *SH3BP1* and *ALDH3A1*, were strongly associated with tau-related pathology and early disease progression (Figure 6B).^41,44^ In contrast, non-overlapping DEGs like *S100A9* and *SIGLEC8* reflected PD-specific changes independent of tau pathology, highlighting distinct molecular pathways (Figure 6B).^43,45^ Late-stage DEGs, including *MEIS3P2* and *CACTIN-AS1*, were linked to glial activation and chronic inflammation, further emphasizing the role of astrocytes in late-stage PD progression (Figure 6B).^46,47^ Similar trends were observed in PDD, where early-stage DEGs like *RHOA-IT1* and *CXCR3* marked tau pathology early disease progression, while non-overlapping genes such as *PSEN2* and *ALDH1A1* highlighted pathways specific to early PDD progression (Figure 6C).^29,49^

This study provides a comprehensive transcriptomic landscape of the CING in PD and PDD, revealing cell-type-specific molecular changes and stage-specific pathways. By distinguishing tau-dependent and independent mechanisms, it identified candidate genes and pathways for therapeutic targeting and biomarker development. Future research should focus on validating these findings in larger cohorts and exploring their functional relevance in disease progression.

### Limitations of the study

It primarily stems from the challenges associated with managing and utilizing human samples, particularly postmortem brain samples. Limited sample availability necessitated concurrent experimentation, which could introduce experimental error, especially when verifying multiple samples and conditions with shallow sequencing. To mitigate this, future studies should focus on optimizing and streamlining sample preparation processes. Additionally, the number of sgRNAs targeting rRNAs or non-specific RNAs via dCas9 may be insufficient. Expanding the variety of target sgRNAs in future experiments could enhance the robustness of results. The inherent constraints of working with postmortem human brain samples also affected data availability, necessitating validation of the DGE analysis results to ensure reliability. The effects of sample size and distribution were carefully analyzed, leading to the identification of a robust gene set minimally influenced by external factors, which serves as candidates for further validation. However, cohort distribution biases across conditions and Braak stages in the PD group could impact the analyses of disease progression. This limitation can be addressed through additional experimental validation or by expanding datasets, guided by the methods and results established in this study. Together, these approaches aim to overcome current limitations and yield more reliable and comprehensive findings.

## Supporting information

All-tablrs

## Author contributions

Conceptualization, J.P., S.U.K., S.H., D.K., B.N., T.M.D. and V.L.D.; Quality control and library preparation, J.P., S.U.K., S.H., D.K., and J.C.T.; Data analysis, J.P., S.U.K., S.H., D.K., Q.P., and B.N.; investigation, J.P., S.U.K., S.H., D.K., Q.P., B.O.V., J.T., S.B., T.M.D., and V.L.D.; resources, S.U.K., S.H., J.R.O., O.P., J.C.T., T.M.D., and V.L.D.; writing—original draft, J.P., S.U.K., S.H., D.K., T.M.D., and V.L.D.; visualization, J.P., S.U.K., S.H., and D.K.; supervision, S.U.K., B.N., T.M.D. and V.L.D.; funding acquisition, S.U.K., T.M.D. and V.L.D.

## Acknowledgements

This work was supported by grants from the National Institutes of Health (NIH)/National Institute of Neurological Disorders and Stroke (NINDS) R01 NS123456 and a sponsored research agreement with Valted Seq. T.M.D. is the Leonard and Madlyn Abramson Professor in neurodegenerative diseases. We thank Hao Wen (Gilbert) Chen and Noelle Burgess, Medical Illustrators at the Johns Hopkins University Institute for Cell Engineering, for creating the illustrations and schematics in this publication. Raw data and meta data for patients can be requested to Ted M. Dawson or Bardia Nezami

## Materials and Methods

### KEY RESOURCES TABLE

| Reagent or Resource | Source | Identifier |
| --- | --- | --- |
| Chemicals |  |  |
| SUPERase RNase inhibitor | ThermoFisher | AM2696 |
| Maxima H- Reverse transcriptase | ThermoFisher | EP0753 |
| Poly(A) polymerase | New England Biolabs | M0276L |
| Non-directional second strand synthesis module | NEW ENGLAND BIOLABS | E6111L |
| Quick Ligation kit | NEW ENGLAND BIOLABS | M2200L |
| DEPC water | Quality Biologicals | 351-068-131CS |
| USER II enzyme | NEW ENGLAND BIOLABS | M5508L |
| dNTP mix | ThermoFisher | R0192 |
| Tagmentase- unloaded | Lucigen | TNP92110 |
| Q5 PCR mix | New England Biolabs | M0491 |
| TDE1 enzyme | Illumina | 20034197 |
| dCas9-3xFLAG-biotin protein | Millipore Sigma | DCAS9PROT |
| NEB next library Quant kit | NEW ENGLAND BIOLABS | E7630 |
| Qiagen protease | Qiagen | 19157 |
| YOYO-1 | Thermo Fisher | Y3601 |
| RNaseZap | Thermo Fisher | AM9780 |
| Agencourt AMPure XP | Beckman Coulter | A63882 |
| Amicon filter | Thermo Fisher | UFC500324 |
| Anti-FLAG M2 magnetic beads | Millipore Sigma | M8823 |
| Protease inhibitor Cocktail tablets | Millipore Sigma | 11697498001 |
| FloMi filter 40µM | VWR | 10032-802 |
| Falcon cell strainer 40µM | VWR | 21008-949 |
| 2% Agarose, PippinHT | Saga science | HTC2010 |
| Sucrose | Millipore Sigma | S0389 |
| TritonX-100 | Millipore Sigma | T8787-100ML |
| Tween 20 | Thermo Fisher | BP-337-100 |
| Magnesium Chloride solution 2M | Millipore Sigma | 68475-100ML-F |
| Sodium Chloride | Millipore Sigma | S3014-500G |
| Potassium Chloride | Millipore Sigma | P9541-500G |
| Igepal CA-630(NP-40) | Millipore Sigma | I8896-50ML |
| DTT | Millipore Sigma | 10197777001 |
| EDTA | Millipore Sigma | E4884 |
| 10X Dulbecco's Phosphate Buffered Saline | Thermo Fisher | 14200075 |
| Tris (1 M), pH 8.0, RNase-free | Invitrogen | AM9855G |
| TAPS buffer (1M, pH 8.5) | Boston Bio | BB-2375 |
| N,N-Dimethylformamide | Millipore Sigma | 227056 |
| DNA binding buffer | Zymo | D4004-1 |
| High Sensitivity D5000 Screen tape | Agilent | 5067-5592 |
| High sensitivity RNA screen tape | Agilent | 5067-5579 |
| Direct zol 96 RNA kits | Zymo research | R2054 |
| Nexteara XT index kit v2 set A | Illumina | FC-131-2001 |
| Nexteara XT index kit v2 set B | Illumina | FC-131-2002 |
| Nexteara XT index kit v2 set C | Illumina | FC-131-2003 |
| Oligonucleotide |  |  |
| SgRNAs | This study | See the Table |
| Software and algorithms |  |  |
| R | N/A | RRID:SCR_001905 |
| Combat-seq | Zhang et al. | 10.1093/nargab/lqaa078 |
| DESeq2 | Love et al. | RRID:SCR_015687 |
| edgeR | Mark et al. | RRID:SCR_012802 |
| Seurat | Hao et al. | RRID:SCR_016341 |
| WebGestalt | Liao et al. | RRID:SCR_006786 |
| KEGG | Minoru et al. | RRID:SCR_012773 |
| ALRA | Linderman et al. | 0.1038/s41467-021-27729-z |

### Shallow bulk RNA seq using the frozen tissue samples

Frozen human tissues (approximately 10 mg) were sectioned on dry ice using a razor blade and homogenized in ice-cold homogenization buffer (250 mM sucrose, 25 mM KCl, 5 mM MgCl2, 10 mM Tris-HCl pH 8, 0.1% RNase inhibitor (RI), 0.1% TX-100, 0.1% IGEPAL CA-630, and DEPC water). Homogenization was performed with 15 strokes of a Dounce homogenizer. The homogenate was filtered through a 40 µm cell strainer and centrifuged at 500 × g for 5 minutes at 4°C. The nuclear pellet was resuspended in cold methanol and fixed at -20°C for 30 minutes. Total RNA was extracted using the Direct-zol 96 RNA Kit according to the manufacturer’s instructions. RNA integrity was assessed by determining the RNA Integrity Number (RIN). For bulk RNA-seq library preparation, 250 ng of total RNA was reverse-transcribed into cDNA using the Maxima H- Reverse Transcriptase kit with an anchored poly-dT primer, following the manufacturer’s protocol. Double-stranded cDNA was synthesized using a non-directional second-strand synthesis enzyme, following the instructions provided.

The double-stranded cDNA was processed for sequencing by tagmentation using the Illumina i5 and i7-adapted Tn5 enzyme. The sample was mixed with i5 and i7 indexed primers and Q5 PCR Master Mix and amplified under the following thermal cycling conditions: initial denaturation at 98°C for 1 minute, followed by 12–14 cycles of 98°C for 10 seconds, 55°C for 30 seconds, and 72°C for 45 seconds, with a final extension at 72°C for 5 minutes. After PCR, the libraries were purified using 0.8× AMPure beads and analyzed using a TapeStation. Subsequently, sequencing was performed on a NextSeq 550 platform, generating approximately 10 million reads per sample with the following cycle settings: Read 1 for 75 cycles, i5 for 8 cycles, i7 for 8 cycles, and Read 2 for 75 cycles.

### Preselection of human post-mortem brain tissue sample using Sample Quality Matrix (SQM)

To identify high-quality samples for snRNA-seq, we performed both bulk RNA-seq and snRNA-sequencing on PD and control samples across multiple brain regions, analyzing a total of 18 samples. Based on snRNA-seq results, we classified samples as high- or low-quality depending on the number of high-quality nuclei obtained. Then we identified over- and under-expressed genes in high-quality samples, referred to as high-quality genes (HQGs) and low-quality genes (LQGs), respectively, compared to low-quality samples based on the corresponding bulk RNA-seq data. We defined SQM (sample quality metric; SQM) as the difference between the average log2 CPM (counts per million) values of high-quality genes (HQGs) and low-quality genes (LQGs). Post-mortem brain tissues from the cingulate cortex (CING) were obtained from two brain banks: 120 samples from the Johns Hopkins University School of Medicine and 120 samples from the Banner Health Institution. Next, a total of 135 patients (45 controls, 45 Parkinson’s disease [PD], and 45 Parkinson’s disease dementia [PDD]) were selected based on the highest SQM as determined by shallow bulk RNA sequencing.

### Processing of HiF-snRNAseq using the frozen CING tissue samples

Approximately 20 mg of the samples were homogenized in cold homogenization buffer with the same composition as the buffer used in Shallow seq using a glass on dounce homogenizer and strokes 15 times softly. The homogenized sample was transferred to the top of a 40 µm cell strainer to eliminate tissue debris and collect the filtrated nuclei.

It was then centrifuged for 5 min at 600 xg, at 4℃. The pellet of nuclei was resuspended and fixed with cold-methanol for 30 min at -20℃. For permeabilization, to the fixed nuclei were added TX-100 for final concentration of 0.1% for 3 min on ice and centrifuged for 5 min at 800 xg at 4℃. The pelleted nuclei were washed once with 50 mM Tris pH 8 with 0.1% RI and spin down at 800 xg for 5 min at 4℃. The pellet was resuspended in 250 µL cold 0.5 X PBS with 1% RI. The sample was passed through 70 µm Flowmi cell strainer. To determine the number of nuclei for Poly-A tailing reaction, the purified sample was stained with DAPI and YOYO-1, and subsequently measured by Countess.

### Enzymatic reaction for Poly-A tailing

Prior to the RT reaction, the following components were added to each sample: 50 units of Poly(A) polymerase (NEB, M0276L), 1 mM ATP, and 1x buffer. 1 unit of enzyme was used per 10 million nuclei. The reaction mixture was then incubated at 37°C for 10 minutes with gentle rotation. To terminate the reaction, 10 mM EDTA was added, and the mixture was centrifuged at 800 xg for 5 minutes, repeated twice.

### 3-step split-pool barcoding

The first barcoding occurs through an *in situ* reverse transcription (RT-PCR). 100K nuclei were aliquoted into each well and added with 100 µM barcoded dT and 10 µM dNTP, denatured at 55℃ for 5 min and placed on ice. The RT mix (1X RT buffer, 0.5U RI, 20U Maxima RT, and DEPC water) was added to each well. The plate was incubated at 4℃ for 2 min, 10℃ for 2 min, 20℃ for 2 min, 30℃ for 2 min, 40℃ for 2 min, 50℃ for 2 min, and 55℃ for 30 min. The samples were pooled together and were centrifuged for 5 min at 800 xg at 4℃. The nuclei pellet was resuspended in PBS and transferred to a new tube.

The second barcoding occurs through a ligation. The pooled nuclei were redistributed into a plate with each well including 100 µM barcoded ligation adaptors, Quick ligation kit (NEB) and incubated at 25℃ for 20 min. After the reaction, samples were pooled together and spun down at 800 xg for 5 min and washed twice with PBS using 800 xg at 4℃ for 5 min. For double strand synthesis, about 5K nuclei were redistributed into well and added with the second strand synthesis buffer and enzyme. Plate incubations were carried out at 16℃ for 180 min and were stopped by 75℃ for 20 min.

For tagmentation, the tag mix was added to each well (1X tn5 buffer, N7 only- Tn5) and incubated at 55°C for 5 min and then stopped by 0.2% SDS buffer at RT for 5min. The DNA was purified using 1X AMPure beads. For elution and USER reaction, to each well was added USER mix (1X USER buffer, 0.1 U USER enzyme and D.W) and incubated at 37℃ for 20 min. The elution sample was transferred into a new plate and incubated at 80℃ for 10 min to stop USER activity.

The third barcoding occurs through a PCR reaction. Each well was mixed with 10 pM indexed i5 and i7 primers and 1X Q5 PCR master mix. Amplification was carried out using the following program: 72℃ for 5 min, 98℃ for 30 s, 15 cycles of (98°C for 10 s, 66°C for 30 s, 72℃ for 45 s) and a final 72℃ for 5 min. The library will be pooled and concentrated using AMPure and purified allowed library size (350 bp to 750 bp) by Pipin HT with 2% gel cassette. Library concentrations were determined by qPCR Library Quantification kit and DNA trace by TapeStation. Libraries were sequenced on NextSeq550 (read 1: 40 cycles, read 2: 100 cycles, index 1: 10 cycles, index 2: 10 cycles) or NovaSeq platform (read 1: 150 cycles, read 2: 150 cycles, index 1: 10 cycles, index 2: 10 cycles).

### dCas9 purification

Target sgRNAs were assembled by IDT, and the manufacturer’s protocol was used to incubate sgRNAs with dCas9-3xFLAG-biotin protein (Millipore Sigma, DCAS9PROT) at room temperature (RT) for 30 min. We then mixed the resulting RNP with PCR products and 1X Cas9 nuclease reaction buffer and incubated them on a rotor at 37°C for 2 hrs. Next, washing FLAG-magnetic beads was performed twice with cold PBS and assembled them with the PCR product and RNP mixture. The mixture was incubated overnight at 4°C and collected the supernatant, which was subsequently concentrated using AMpure beads.

### Data processing and quality control for read analysis

The raw sequencing files (bcl) were demultiplexed using Illumina’s bcl2fastq (v1.8.4) tool producing raw sequencing reads (fastq). These raw sequencing reads were processed through a customized pipeline for read analysis. Cutadapt (v4.0) was used to identify and trim the ligation barcodes, linker sequence, unique molecular identifier (UMI), RT barcodes and poly-T sequences.^64^ The reads with all the barcodes were further cleaned and trimmed for low quality using BBduk (v38.95) (https://jgi.doe.gov/data-and-tools). The reads were then aligned to Homo sapiens reference genome (genome-version: GRCh38, genome-build: GRCh38.p13) using STAR (v2.7.2a).^65^ Unique mRNA molecules which were amplified during library preparation were identified using UMI-tools dedup method (v.1.1.4).^66^ The final quantification of deduplicated reads was performed using a custom script to account for exonic, intronic, intergenic and reverse reads. Cells with >200 UMIs, <5000 UMIs, >50 genes and < 20% MT genes were used for analysis. Additionally, samples with > 3000 cells were used, and the number of samples remaining was 77.

SCTransform was applied for the raw count matrix normalization and variance stabilization of molecular expression count. We then applied an adaptively-thresholded low rank approximation (ALRA) algorithm for imputing the missing values of the count matrix. Next, we performed principal component analysis (PCA) by using the 3,000 highly variable genes (HVGs) then uniform manifold approximation and projection (UMAP) analysis for lower dimensional projection (number of PCs = 50). For more accurate cell type annotation, we applied transfer learning strategy by using four reference brain snRNA seq datasets with different technology (SMARTseq, 10X, and split-seq) and different brain regions (sACC, hippocampus, multiple cortical area, and temporal gyrus). We identified cell type for each cell using the following procedure: 1. Performing UMAP reduction analysis for a given sample dataset. 2. Performing UMAP reduction analysis for a given reference dataset. 3. Finding common transfer anchor pairs (an anchor pair is a pair of cells each from the sample and reference dataset that are mutually nearest neighbors) using the genes that are common in both the reference and sample dataset. 4. Transferring cell type information of the reference dataset into the sample dataset and projecting the sample’s low-dimensional embeddings into the reference’s low-dimensional embeddings. For each sample, we applied the upper- mentioned steps across four reference datasets and six PC dimensions used for projection. To obtain the optimal annotation among the combination of four reference sources and six PC dimensions, the Silhouette coefficient was used as a clustering evaluation metric.^67^ The Silhouette coefficient is a metric used to calculate the goodness of a clustering technique considering intra-cluster distance and inter-cluster distance.

After averaging the silhouette coefficient of all samples, the combination with the highest value among the various combinations was selected for further analysis.

### Differential gene expression analysis based on pseudo-bulk gene expression profiles

Raw read counts at the single-cell level were transformed into sample-level data per cell type to perform pseudobulk differential gene expression (DGE) analysis.^68^ To correct for batch effects between sample batches, ComBat-seq was applied to the pseudobulk data and then DGE analysis was performed separately for each cell type using edgeR.^69,70^ Before performing downstream analysis, a chi-square test to determine if there were statistically significant differences in the number of cells across conditions by cell types (PMID: 23894860, 39850354). Lowly expressed genes, which are genes with a total expression level of less than 10, were then filtered out and outlier samples were excluded based on the PCA mapping results. Wald tests were performed on three contrasts (PD versus control, PDD versus control, and PDD versus PD) to define differentially expressed genes (DEGs) with adjusted p-values (FDR) less than 0.05.

Since the low-dimensional mapping results are different for each cell type and thus the outliers that are removed are different, the number of samples are different when performing DGE analysis for each cell type.^71^ By comparing the DEGs obtained for these three contrasts, genes were identified that are unique to each contrast and genes with shared characteristics. Additionally, to compare the identified DEGs with already known pathways, pathway analysis was conducted using the WebGestalt for KEGG pathway.^72,73^ To compare single-cell based analysis with those of shallow bulk sequencing, the same approach was applied to the shallow bulk RNA sequencing data. DE analysis was performed using the same samples as the single cell study, considering disease progression of PD and PDD, respectively to examine disease progression. Using the obtained DEGs, correlated genes that showed same trends between single-cell and shallow bulk studies using fold-change values were examined.

### Robustness and stability analysis using random subsets

To verify the reliability of the DGE analysis results using our samples, analyses using random subsets were performed to identify the impact of sample size on the DE analysis results and variance between individuals within the same conditions. Random sampling was performed using the minimum number of samples for the three conditions as a baseline. In other words, random sampling without replacement was performed by dividing the entire cohort into random subsets based on the minimum number of each condition. To determine the number of iterations at which the results sufficiently converge, an experiment with different number of iterations was performed considering the mean and median values. The similarity between the mean and median was used as statistical stability, and it was confirmed that our results converge after 1000 iterations. Consequently, random sampling was iterated 1000 times, and DGE analysis was performed using each randomly sampled subset. The number of overlapping genes between DEGs obtained from each iteration and DEGs obtained using the entire sample was examined for each contrast and cell type. We investigated these overlapping genes from two aspects. The first overlap ratio (Ratio1) is calculated by dividing the number of overlapping genes by the number of DEGs obtained using the entire sample and then averaging over iterations. This Ratio1 indicates how the number of samples affects the DE analysis results. The second overlap ratio (Ratio2) is calculated by dividing the number of overlapping genes by the number of DEGs obtained per random sampling iteration then also averaging over iterations, so this Ratio2 represents the variances between individuals within the same conditions.

### Examination of PD and PDD progression considering Braak stage

To determine how individual cell types are affected by clinical variables, DGE analysis was performed for each cell type considering clinical variables of age, Braak stage, and gender.^70^ To calculate the likelihood of dispersion estimation, the design matrix for each variable and its pair was used to perform DGE analysis. Using interaction models, we performed analyzes that considered not only how each variable affected the conditions, but also how their combined effects differed from those predicted based on their individual effects.

To investigate the progression of PD and PDD, DGE analysis according to disease progression was performed using the Braak stage, which was the variable that has the greatest influence on the DGE analysis. Based on the clinical information, we categorized Braak stage I and II as early stage and stages III, IV, and V as late stage. To investigate how cell type composition influences disease progression, we compared the number of cells across Braak stages for all cell types and examined cell abundance along disease progression. Then, for each cell type, DGE analysis was performed considering disease progression in PD and PDD, respectively. Since there was Braak information based on pathological diagnosis for control samples, DGE analysis was performed using control samples stratified into early- and late-stages. DGE analysis was performed using the same method as before (Method: Differential gene expression analysis). By comparing the DEGs obtained considering disease progression, early- or late-specific genes were identified. Additionally, DGE analysis considering disease progression using all control samples as control (unstratified) was performed to figure out the consistent markers of disease progression that are independent of the stratification strategy.

### In-depth analysis of disease progression and Braak stage

To in-depth examine DEGs considering disease progression, DGE analysis was performed considering only the Braak stages regardless of conditions. For each cell type, Wald test was performed comparing Braak-early and Braak-late samples to define DEGs only related to Braak stages independent of condition (control, PD, and PDD) with FDR less than 0.05. After obtaining DEGs considering only the Braak stages, they were divided into up- and down-regulated genes. The baseline condition of the Wald test was Braak-early, so up-regulated refers to late-related and down-regulated refers to early-related. Using obtained DEGs, stage-specific DEGs considering disease progression were break into Braak-dependent and independent.

---

**Figure S1.**
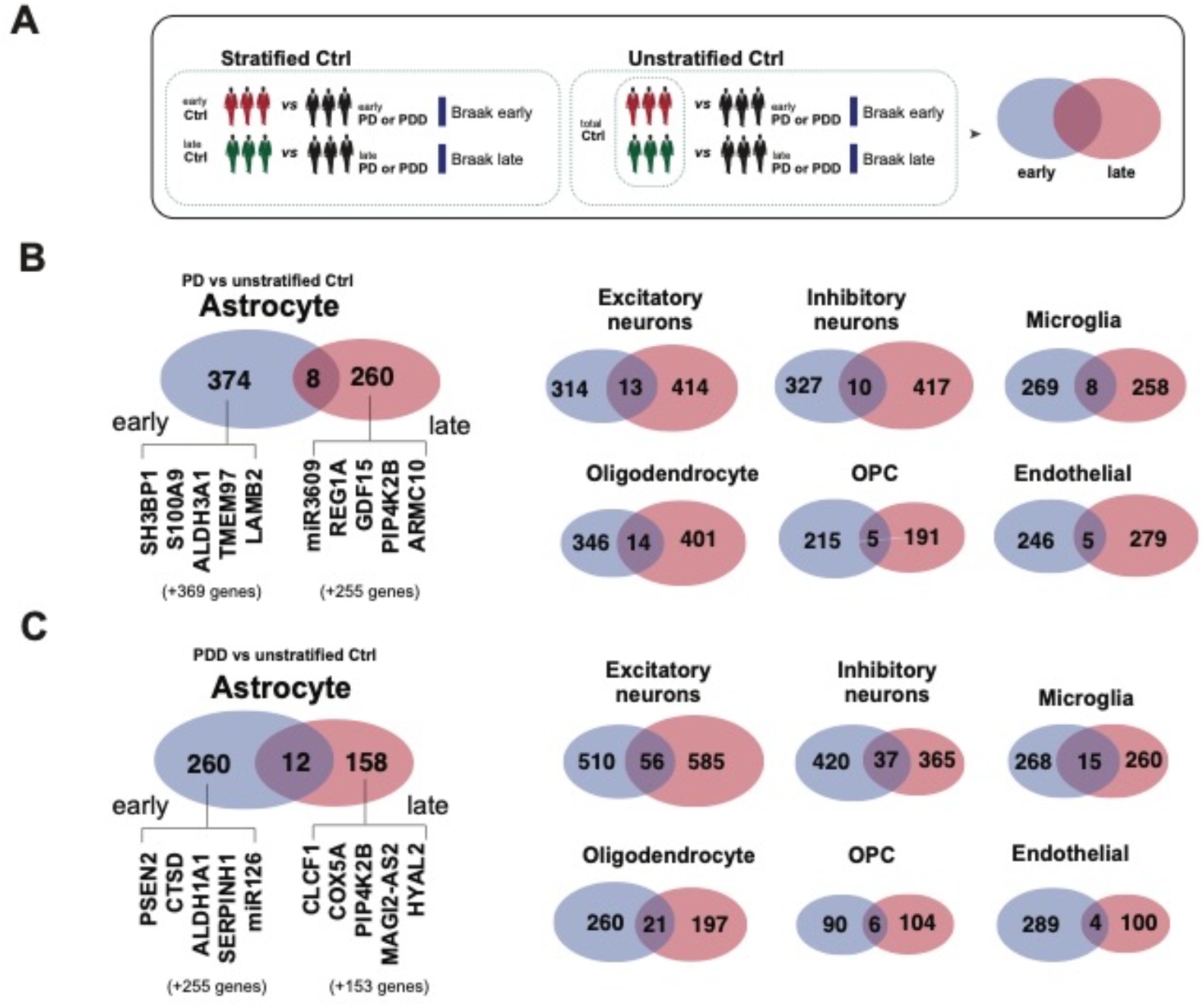
Comparison of differentially expressed genes using unstratified control (control) and Braak-stratified PD and PDD groups. (A) Diagram for the analytical approach: stratified controls and unstratified controls, (B) Venn diagrams for PD versus control showing early- and late-stage DEGs with unstratified controls in astrocytes with the representative DEGs and other major cell types, and (C) DEGs from the comparisons of PDD versus control.

**Figure S2.**
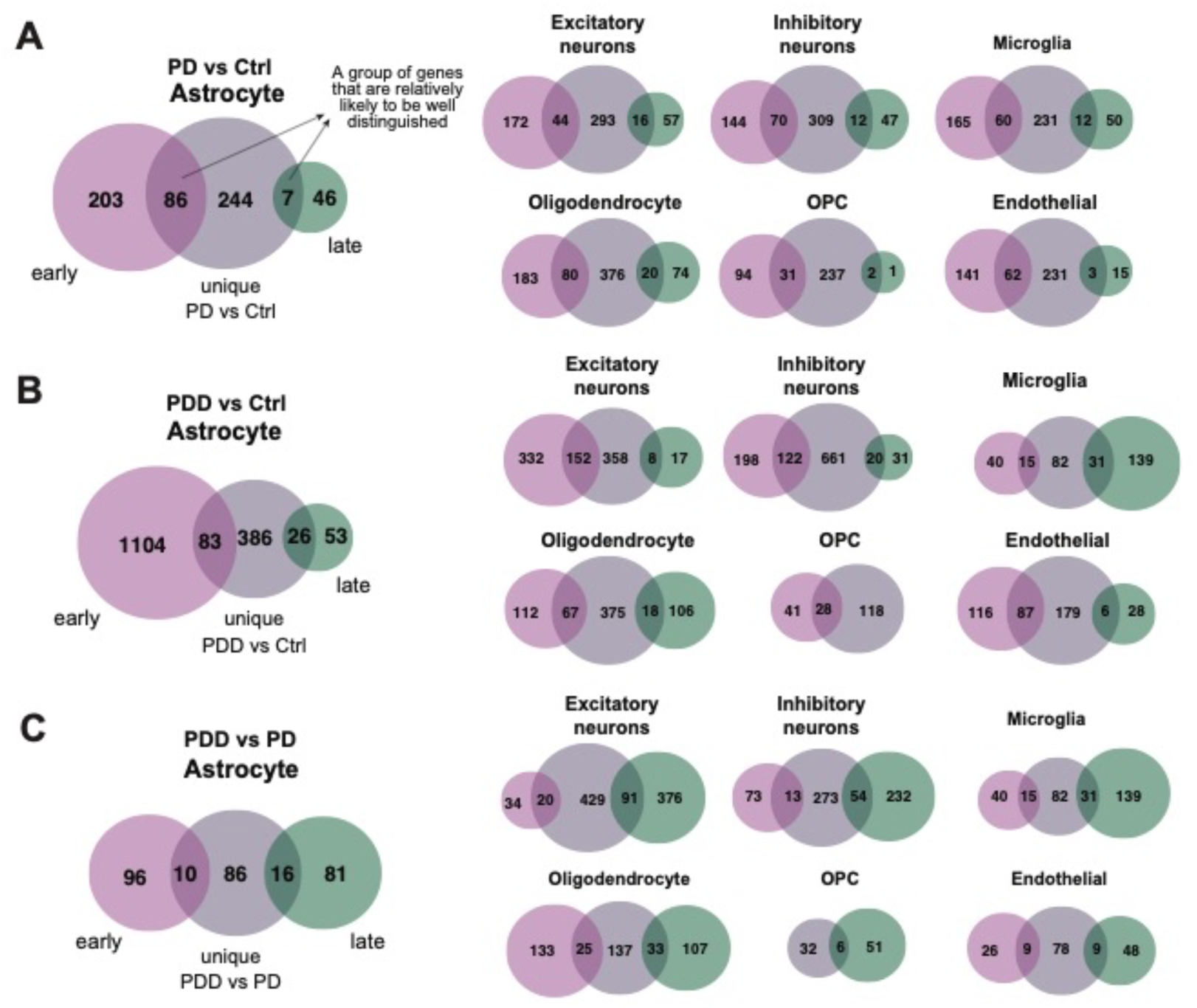
Comparison of DEGs across contrast-unique and stage-specific disease conditions. (A) Venn diagrams comparing early- and late-stage DEGs for PD versus control in astrocytes and other major cell types using contrast-unique and stage-specific, (B) for PDD versus control, and (C) for PDD versus PD.

**Figure S3.**
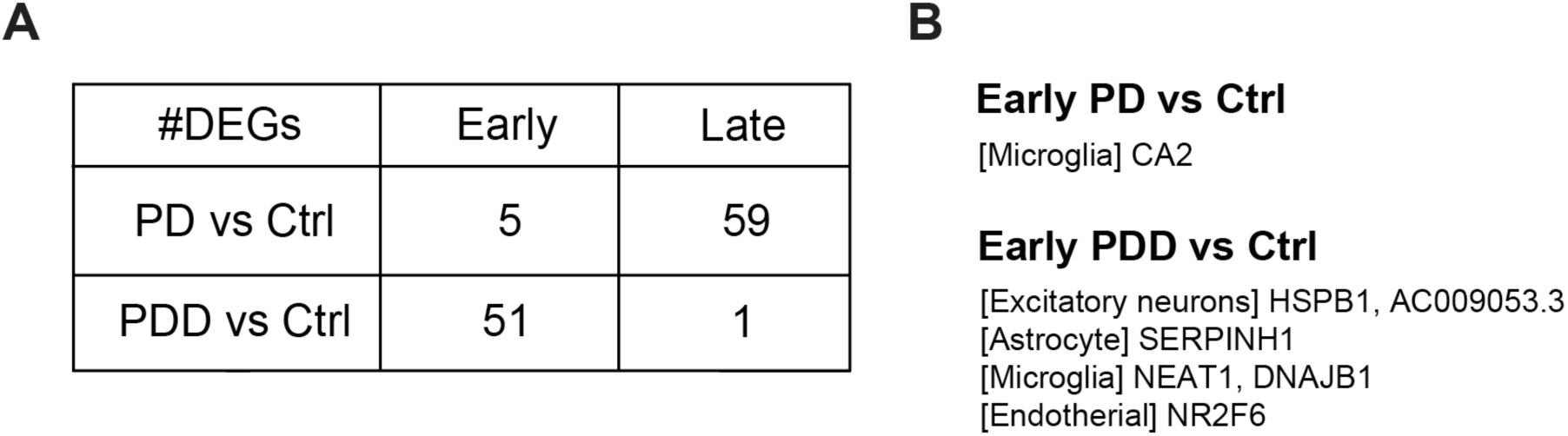
Stage-specific differentially expressed genes across PD and PDD conditions using shallow bulk RNA sequencing. (A) Summary of DEGs in early (stages I-II) and late (stages III-V) Braak stages for PD versus control and PDD versus control. (B) Genes obtained from comparisons of early-stage DEGs identified in PD versus control and PDD versus control across DEGs obtained from HiF-snRNA-seq.

**Table S1.** Demographic and clinical characteristics of the selected patient cohort with RIN and SQM values

**Table S2.** List of dCas9-targeted gRNA for optimizing snRNA-seq from post-mortem brain tissue

**Table S3.** Gene count matrix for 2.0 M nuclei from human cingulate cortex

**Table S4.** Condition-specific DEGs across seven major brain cell types

**Table S5.** KEGG pathway enrichment analysis of DEGs by cell type and condition

**Table S6.** Gene list resulted from the comparison of early and late Braak stages in stratified control, PD, and PDD groups

**Table S7.** Gene list resulted from the comparison of early and late Braak stages in unstratified control, PD, and PDD groups

**Table S8.** Gene list from the comparisons of experimental contrasts and Braak stages

**Table S9.** Gene list resulted from shallow bulk RNA sequencing with the comparisons of early and late Braak stage between control and PD or PDD

**Table S10.** Stage-specific DEGs Linked to Tau-dependent and independent mechanisms

